# Age-related Reorganization of Corticomuscular Connectivity During Locomotor Perturbations

**DOI:** 10.1101/2025.09.28.679054

**Authors:** Seyed Yahya Shirazi, Shahamat Mustavi Tasin, Helen J. Huang

## Abstract

Locomotor perturbations elicit cortical and muscular responses that help minimize motor errors through neural processes involving multiple brain regions. The anterior cingulate cortex monitors motor errors, the supplementary motor areas integrate sensory and executive control, and the posterior parietal cortices process sensorimotor predictions, while muscles show increased activation and co-contraction patterns. With aging, these neural control strategies shift; older adults demonstrate less flexible cortical and muscular responses, using compensatory overactivation and simpler muscle synergies to maintain performance comparable to young adults. We investigated corticomuscular connectivity patterns during perturbed recumbent stepping in seventeen young adults (age 25±4.9 years) and eleven older adults (age 68±3.6 years) using high-density EEG (128 electrodes) and EMG from six bilateral muscles. Brief mechanical perturbations (200ms of increased resistance) were applied at left or right leg extension-onset or mid-extension during continuous stepping at 60 steps per minute. We applied independent component analysis, source localization, and direct directed transfer function to quantify bidirectional information flow between cortical clusters and muscles in theta (3-8 Hz), alpha (8-13 Hz), and beta (13-35 Hz) bands. Young adults demonstrated concentrated electrocortical sources in anterior cingulate cortex, bilateral supplementary motor areas, and bilateral posterior parietal cortices, with strong theta-band synchronization following perturbations. In contrast, older adults showed fewer differentiated cortical sources, particularly lacking distinct anterior cingulate activity, and exhibited only minimal synchronization changes. Baseline corticomuscular connectivity was significantly stronger in older adults compared to young adults (p=0.012), suggesting fundamental differences in resting motor control states. During perturbations, young adults employed flexible, task-specific connectivity modulation involving error-processing networks, with the anterior cingulate showing selective bidirectional connectivity changes with specific muscles. Older adults relied on more diffuse (i.e., not focused to specific brain area) connectivity patterns dominated by motor and posterior parietal cortices, with strong connections to multiple upper and lower limb muscles simultaneously. These findings reveal an age-related strategic reorganization from dynamic, error-driven neural control to a more constrained, stability-focused approach that may reflect compensation for sensorimotor changes. The distinct connectivity signatures establish perturbed recumbent stepping as a valuable tool for assessing corticomuscular communication and provide normative benchmarks for developing targeted rehabilitation interventions to restore efficient motor control in aging and neurological populations.

## 1 Introduction

Cortical and muscular activities are often evident in responses to motor and locomotor perturbations, which may reflect neural processes involved in attenuating motor errors. Several cortical and subcortical brain areas are known to monitor motor errors. The anterior cingulate cortex in the mid-prefrontal brain region is likely to use sensory inputs to detect motor errors relative to the expected goal (Holroyd & Coles, 2002). The supplementary motor area integrates neural pathways from the motor cortex and anterior cingulate and likely acts as a sensory and executive hub for processing errors (Peterson & Ferris, 2019). The posterior parietal cortex is thought to process the differences between the visual and sensorimotor inputs (Peterson & Ferris, 2018). Nevertheless, the activities of these brain areas are not exclusive to these roles and processes. For example, the left posterior parietal cortex has a critical role in joint impedance control, while the right prefrontal cortex controls the limb speed during perturbed upper limb motor tasks (Mutha et al., 2014). Muscles tend to show increased activation and agonist/antagonist co-contraction in response to perturbations (Huang & Ahmed, 2014b; Thoroughman & Shadmehr, 1999). Not surprisingly, corticocortical, intramuscular, and corticomuscular coherence also increase with perturbations indicating greater cortical control over muscular activation (Gentili et al., 2015; Sato & Choi, 2019; Zandvoort et al., 2019).

With increasing age, cortical and muscular activity tend to be less flexible and demonstrate compensatory changes. Older adults often use more co-contraction in response to motor perturbations and may not reduce their muscle activity as they gain more experience with the perturbations as much as young adults (Huang & Ahmed, 2014a). However, locomotor perturbation studies have indicated that older adults use simpler muscular control through fewer muscle synergies or fewer driving muscles to maintain similar motor performance as young adults (Allen & Franz, 2018; Da Silva Costa et al., 2020). Many studies have shown “overactivation” of older adults’ brain regions compared to young adults as a compensatory mechanism for similar motor performance (Reuter-Lorenz & Cappell, 2008; Seidler et al., 2010). Such overactivation would increase connectivity at both cortical and muscular levels when facing motor challenges (Johnson & Shinohara, 2012; Kamp et al., 2013; Walker et al., 2020). In contrast, a couple of studies on standing and locomotor tasks using channel electroencephalography (EEG) and electromyography (EMG) signals indicated weaker connectivity in older adults (Ozdemir et al., 2018; Roeder et al., 2020; Yoshida et al., 2017). The different methods for computing connectivity, lack of source-resolved EEG, and inter-subject variability may contribute to the different outcomes observed in the connectivity studies (Clark et al., 2020; Höller et al., 2017).

In individuals with neurological impairments, decreased corticomuscular connectivity is well-documented (Chen et al., 2018; Larsen et al., 2017; Yokoyama et al., 2020), with rehabilitation often improving connectivity or using it as a metric of motor recovery (Chowdhury et al., 2020; Youssofzadeh et al., 2016). We previously demonstrated that perturbations during recumbent stepping elicit brain dynamics similar to perturbed walking, particularly in the anterior cingulate cortex and supplementary motor areas (Shirazi & Huang, 2021). We also showed that older adults recruited fewer muscle-pairs than young adults during perturbed recumbent stepping, despite having similar motor errors as young adults (Shirazi & Huang, 2024). The main advantage of recumbent stepping and other seated exercises is that subjects would not require to maintain their balance while performing the exercise, which is especially beneficial for accessible gait rehabilitation. Establishing connectivity markers during perturbed recumbent stepping provides baseline metrics for rehabilitation applications targeting corticomuscular communication.

The purpose of this study was to investigate corticomuscular connectivity of young and older adults in response to brief perturbations during a locomotor task to identify potential age-related differences in corticomuscular connectivity despite similar behavioral responses. We also aimed to determine whether the timing of the perturbations modulated corticomuscular connectivity. We hypothesized that the corticomuscular connectivity would increase around the perturbation timing, especially between the motor cortex and the driving muscles, namely, posterior deltoids and rectus femoris. This hypothesis was based on our previous EEG and EMG analyses showing increased electrocortical and muscular activation around the stepping perturbation (Shirazi & Huang, 2021, 2024). We also hypothesized that the anterior cingulate cortex would have increased direct causal connectivity from a subset of muscles around the perturbation timing, which would act as sensory feedback for error monitoring. In this study, we use the terms connectivity and causal relation interchangeably.

## 2 Methods

Seventeen young adults (11 females, age 25 *±* 4.9 years) and 11 older adults (4 females, age 68 *±* 3.6 years) participated in the study. Subjects were all right-handed, as determined by their preference for picking up objects, and reported no neurological or physical problems in the two years prior to testing. The Institutional Review Board of the University of Central Florida approved the study, and subjects gave their written informed consent before starting the experiment.

### 2.1 Hardware

We used a recumbent stepper integrated with a servomotor (Shirazi & Huang, 2019) to introduce brief perturbations in the form of added resistance during stepping. The stepper was mechanically coupled such that the contralateral arm and leg extended together. We used the servomotor to briefly increase stepping resistance for 200 ms as the subject’s left or right leg was at the extension-onset or mid-extension. The increased stepping resistance was a torque-controlled algorithm that would require 3x the normal torque to drive the stepper at 60 steps per minute. We used the servomotor’s position sensor to record the stepper’s kinematics at 100 Hz.

We used twelve wireless electromyography (EMG) sensors (Trigno, Delsys, Natick, MA) to collect muscular activity at *∼*1.1 kHz. We attached the EMG sensors on the tibialis anterior, soleus, rectus femoris, semitendinosus, anterior deltoid, and posterior deltoid on both left and right sides (Hermens et al., 2000). For connectivity analyses, we focused exclusively on left-side muscles to examine non-dominant limb control (all participants were right-handed), as this may elicit more pronounced cortical activity. Additionally, preliminary analyses indicated that including all muscles decreased connectivity estimation accuracy due to computational constraints of the multivariate autoregressive model. We used high-density electroencephalography (EEG) system (ActiveTwo, 128 electrodes, BioSemi B.V., Amsterdam, the Netherlands) to collect brain activities at 512 Hz. We placed the EEG cap and electrodes according to the BioSemi manual and used a 3D scanner (Structure I, Occipital Inc., CO) to record the electrode’s 3D locations (Akalin Acar & Makeig, 2013). Data streams of the different systems were synchronized using a trigger signal from the stepper controller to the EMG and EEG systems.

### 2.2 Protocol

Data collection consisted of four 10-minute stepping tasks, each including only one perturbation type at the mid-extension or extension-onset of the left or right legs. Each task started and ended with two-minutes pre and post unperturbed stepping blocks and included a six-minute-long perturbation block, in which subjects would experience perturbation at each stride. There was also a random one-in-five catch stride in the perturbation block, which did not include perturbations. Before starting each task, we instructed subjects to: 1) follow the visual pacing cue presented at 60 steps per minute, 2) step smoothly as if walking, and 3) use both arms and legs to drive the stepper. A complete explanation of the protocol has been presented previously (Shirazi & Huang, 2021). We determined the stepping events using the kinematic information from the stepper servomotor. Each stride was from the start of the perturbed-leg extension to the next perturbed-leg extension. Perturbation events were the start of the increased stepping resistance for each stride.

### 2.3 EEG Processing

EEG and EMG data were processed and analyzed in MATLAB environment (R2018b, MathWorks Inc., Natick, MA) with a custom pipeline based on EEGLAB version 2019.0 (Delorme & Makeig, 2004) (Figure 1). The EEG pre-processing pipeline is largely similar to our previous study on young adults’ perturbed stepping (Shirazi & Huang, 2021). In summary, we used a high-pass filter at 1 Hz and a 60 Hz line-noise filter to clean the EEG data before applying the template correlation rejection method to identify the channels that had a high correlation to the stepping events (Nordin et al., 2019). We then used step-wise channel and frame rejection to create 32 datasets for each subject with a range of conservative to lenient noise rejection. The signal noise rejection metrics were range, variance, kurtosis, and correlation to other channels (Delorme et al., 2013). The frame noise rejection metric was the EEG inter-quartile variability to the overall median of the signal during each task. We used adaptive mixture independent component analysis (AMICA) to quantify the independent components (ICs) for each of the 32 datasets, estimated the IC locations in the brain using DIPFIT, and used a multivariate classifier (ICLabel) to estimate the number of “brain” ICs for each dataset (Palmer et al., 2012; Pion-Tonachini et al., 2019). We ultimately used the dataset with the most “brain” ICs as the representative dataset for each subject.

**Figure 1.**
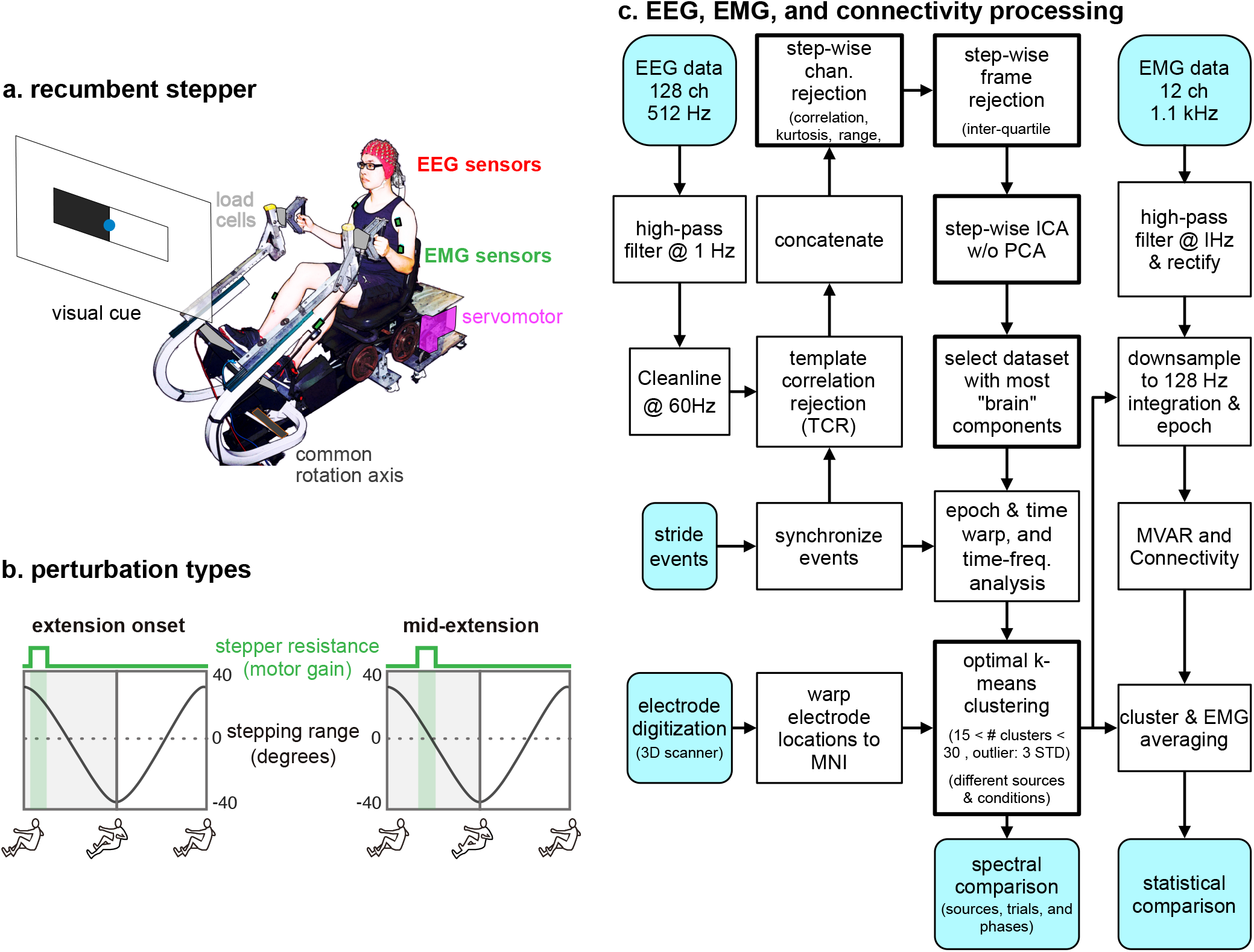
(a) The experimental setup showing the recumbent stepper, pacing visual cue, and EEG and EMG sensors. (b) Shaded green areas show when the perturbations occurred within the stride cycle. (c) EEG, EMG, and connectivity processing pipelines. The thick outlined boxes indicate novel methods developed in this research. The light blue shaded boxes indicate inputs or outputs.

We clustered ICs from young and older adults separately based on the IC location, power spectrum, and scalp map. We used the 3 to 25 Hz range for power spectrum and the Laplacian of the scalp map to perform the clustering (Gramann et al., 2018). We used optimal k-means clustering to select the number of cortical clusters for young and older adults separately and focused on the clusters that included *>*70% of the subjects. If the ICs in a cluster were mostly in a Brodmann area, we would assign that cluster with that Brodmann area (Akalin Acar & Makeig, 2013; Brodmann, 1909). If a cluster spanned across multiple Brodmann areas, we would attribute the cluster with the region that would encompass the cluster’s ICs.

Finally, we computed the temporal profile of the spectral power of each cluster, commonly known as the event-related spectral perturbations (ERSP) (Makeig et al., 2004). We epoched the EEG data from -400 ms to +400 ms of the perturbation events. We baseline normalized the spectral power using the average spectral power during pre and computed the statistically significant event-related synchronization and desynchronization for the ERSPs using EEGLAB permutation functions with alpha equal 0.05 (Delorme et al., 2011). ERSP images only show significant spectral fluctuations.

### 2.4 Connectivity Processing

We used cortical source clusters and EMG from six upper and lower limb muscles to compute connectivity metrics using direct directed transfer function (dDTF) (Korzeniewska et al., 2003; Peterson & Ferris, 2019) (Figure 1). We selected each subject’s ICs that contributed to the cortical clusters and added the EMG signals after high-pass filtering at 1 Hz and rectifying (Myers et al., 2003). We epoched the integrated IC and EMG data from -2 sec to +2 sec of the perturbation events to provide the required data for estimating the multivariate autoregressive models for each subject. We used the Viera-Morf algorithm with 1.3-sec window size and 0.01-sec step-size window to fit a multivariate autoregressive model with an order equal to 32 to each subject’s integrated dataset (Korzeniewska et al., 2003; Vieira & Morf, 1978). We used the 1.3-sec window size to improve the model fit and the 0.01-sec window step-size to provide 10 ms temporal resolution for the connectivity analysis. The model order of 32 provides a spectral resolution of 4 Hz for the connectivity analysis and lets the autoregressive model take more of the past data into the model. The higher model orders, up to 40, have not shown significant noise increases inherent to estimating a greater number of variables in a similar study (Peterson & Ferris, 2019).

### 2.5 Statistical Analysis

We performed a 2×2×2×2 mixed-design repeated measures ANOVA to examine the effects of perturbation side (left/right), task timing (extension-onset/mid-extension), and connectivity direction (brain-to-muscle/muscle-to-brain), with age group (young/old) as the between-subjects factor. The analysis included brain-muscle connectivity from 15 young and 10 older adults. Mauchly’s test confirmed sphericity for all within-subject factors (W=1.0). Box’s M test indicated homogeneity of covariance matrices (p=0.076). When we detected significant interactions, we conducted post-hoc pairwise comparisons using Bonferroni correction to control for multiple comparisons. We set statistical significance at p*<*0.05, with marginal effects reported at p*<*0.10.

## 3 Results

Young adults’ EEG resulted in more individual “brain” ICs and more group-level clusters than older adults (Figure 2). The median “brain” component for young adults was 22 per subject, which was significantly greater than the 10 per subject median for older adults (Wilcoxon rank-sum test, p*<*0.0005). Young adults had five clusters with ICs from *>*70% of the subjects. The clusters were at the anterior cingulate cortex, right supplementary motor area, left supplementary motor area, right posterior parietal cortex, and left posterior parietal cortex. Older adults had only three clusters with ICs from *>*70% of the subjects. Since the ICs within the clusters were more spread to assign a Brodmann area to the clusters, only the general cortical area was attributed to the older adult clusters. The clusters were at the motor cortex, left posterior parietal cortex, and right posterior parietal cortex.

**Figure 2.**
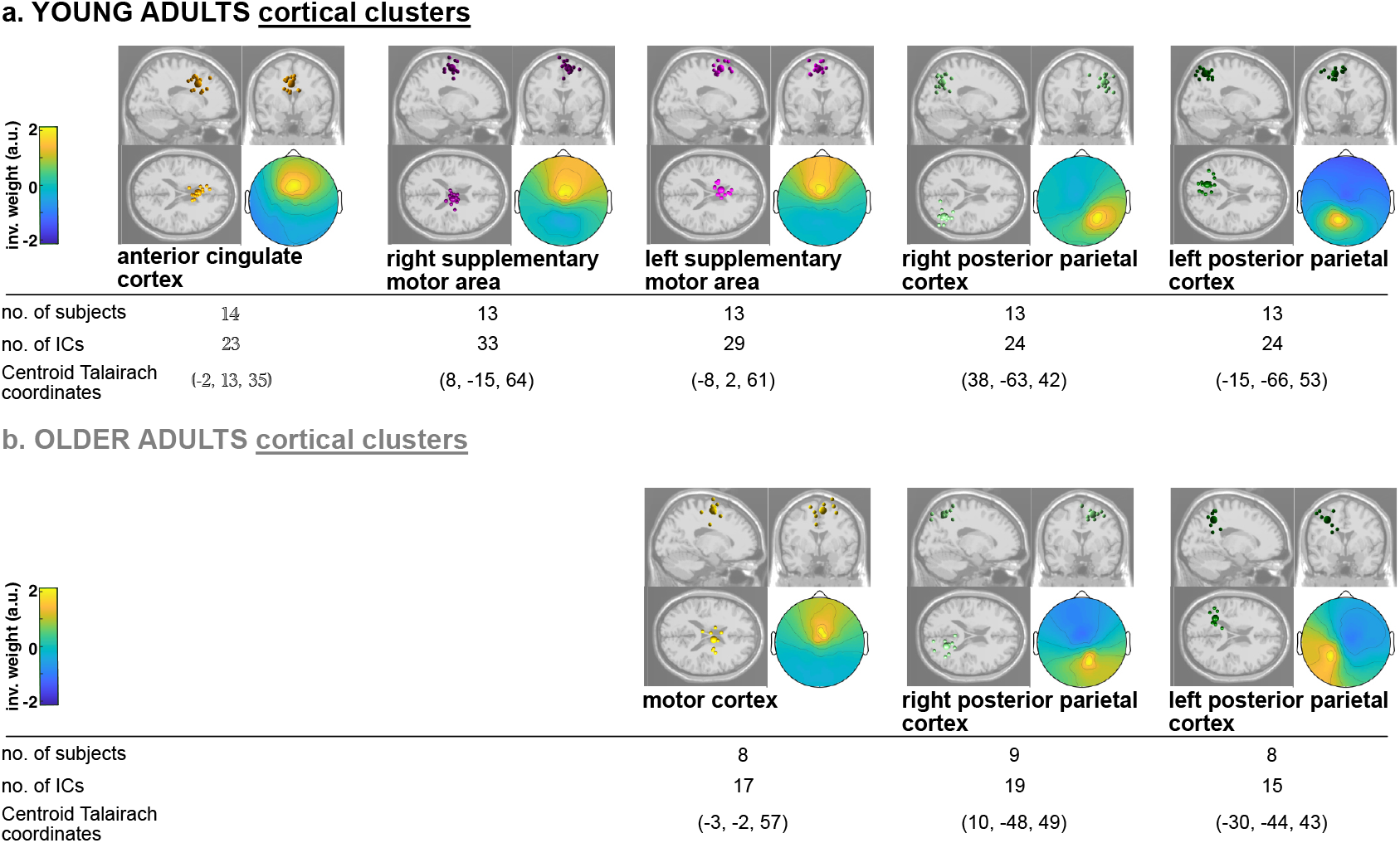
The location of the group cortical clusters for young and older adults. Older adults did not have an anterior cingulate cortex or differentiated motor cortex cluster like the young adults. Both young and older adults had left and right posterior parietal clusters.

### 3.1 Connectivity of Young Adults

Young adults had strong theta-band (3-8 Hz) synchronization locked to the perturbations in the anterior cingulate and supplementary motor areas during left and right extension-onset. However, older adults only showed slight theta synchronization after the perturbations in the motor cortex during left extension-onset (Figures 3, 4). The left posterior parietal cortex had increased theta and alpha (8-13 Hz) synchronization for both young and older adults after the perturbation event. However, the right posterior parietal cortex did not show significant spectral fluctuations locked to the perturbation in either young and older adults (Figures 3, 4). Both young and older adults also demonstrated theta-alpha desynchronization before the perturbation event in the left posterior parietal cortex.

**Figure 3.**
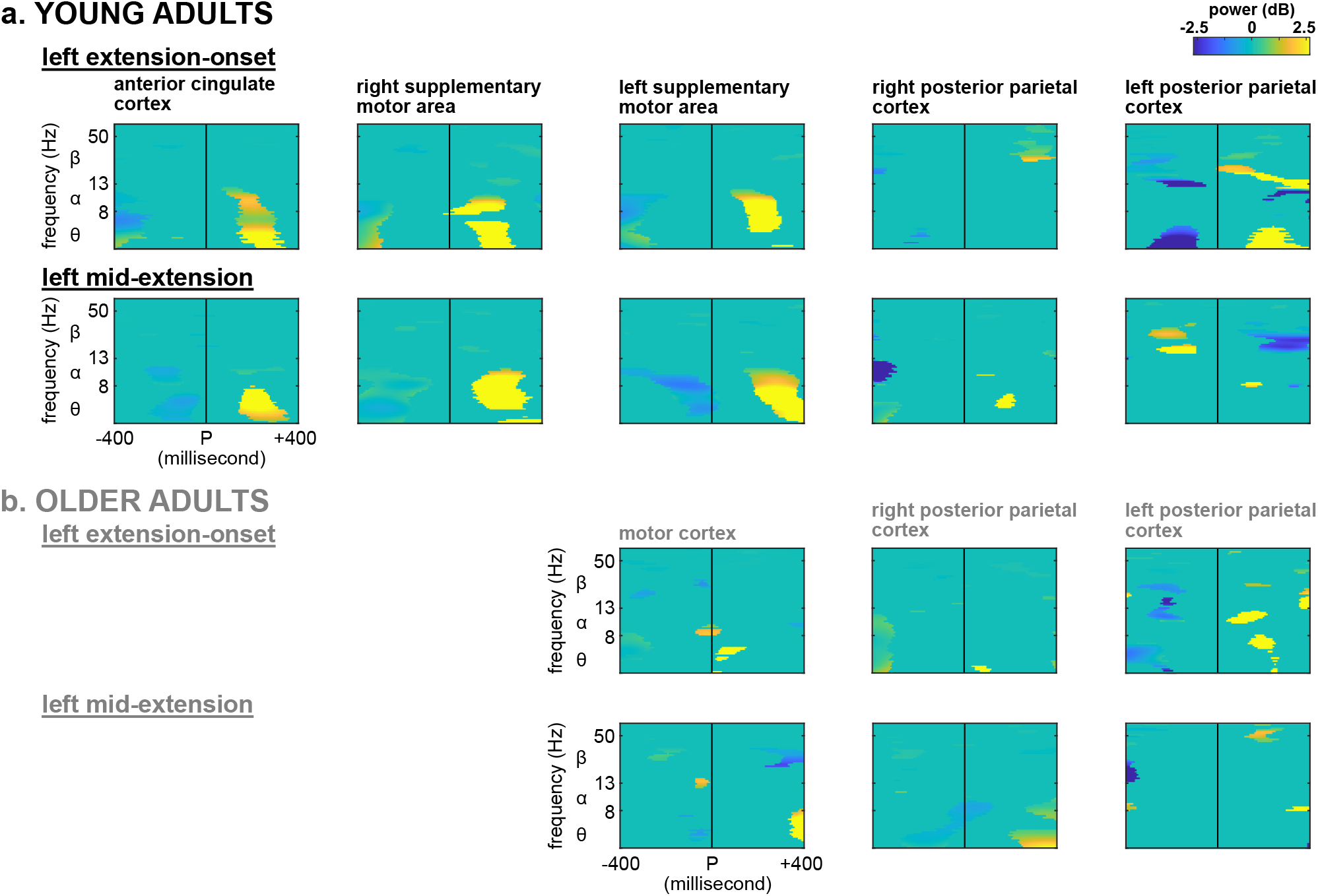
Left extension-onset and left mid-extension event related spectral fluctuations (ERSP) around the perturbation event (P) for young and older adults. Young adults’ anterior cingulate cortex, supplementary motor areas, and left posterior parietal cortex had significant theta-band (3-8 Hz) synchronization after the perturbation event. Older adults had minor theta (3-8 Hz) and alpha (8-13 Hz) synchronization in their motor cortex and left posterior parietal cortex. Yellow areas show greater synchronization while blue areas show greater desynchronization.

**Figure 4.**
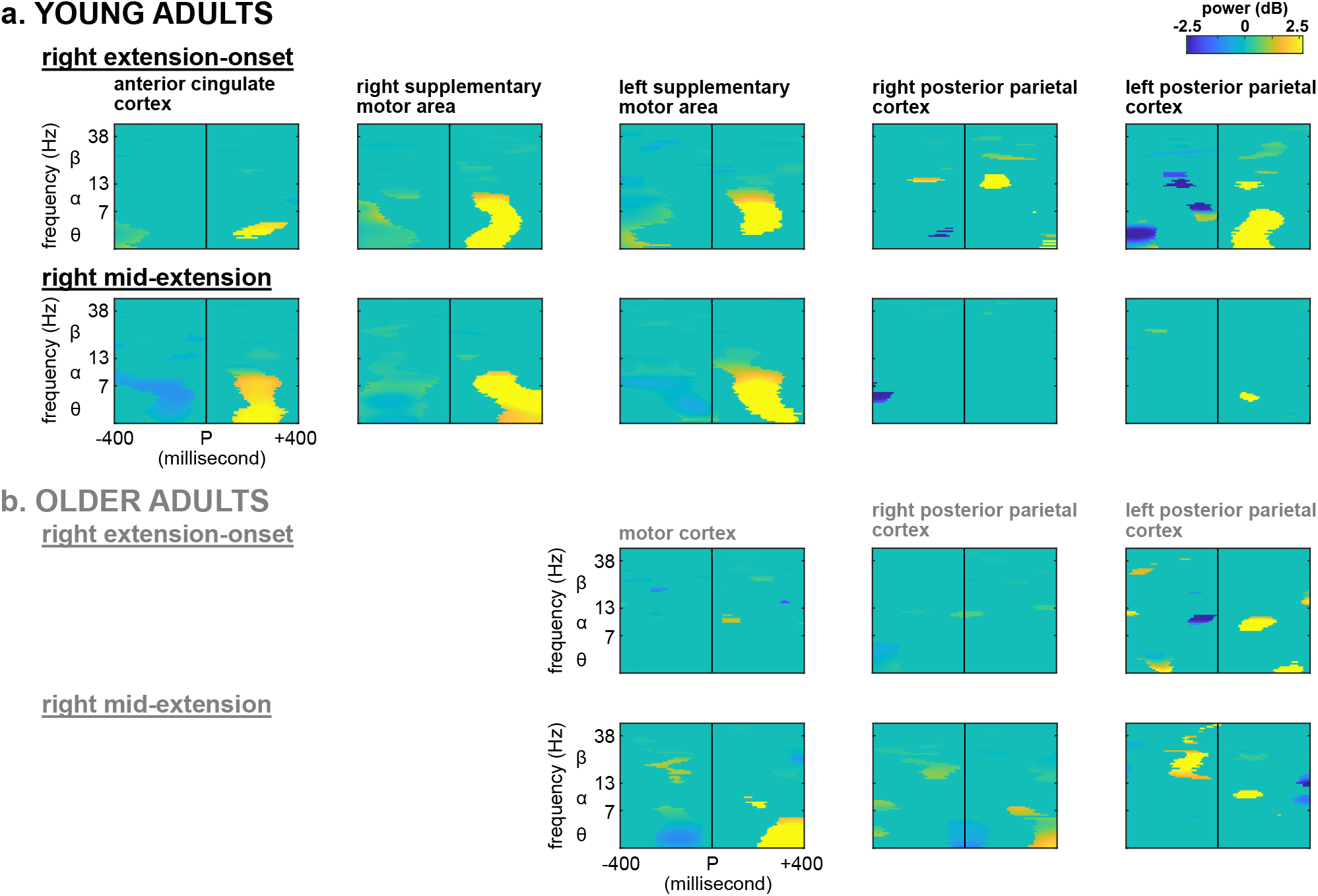
Right extension-onset and right mid-extension event related spectral fluctuations (ERSP) around the perturbation event (P) for young and older adults. Young adults’ anterior cingulate cortex, supplementary motor areas, and left posterior parietal cortex had significant theta-band (3-8 Hz) synchronization after the perturbation event. Older adults had theta (3-8 Hz) and alpha (8-13 Hz) synchronization in the left posterior parietal cortex during extension-onset, and theta synchronization in the motor cortex during mid-extension. Yellow areas show greater synchronization while blue areas show greater desynchronization.

Mid-extension perturbations also elicited theta-band synchronization in the young adults’ anterior cingulate and right and left supplementary motor areas (Figures 3, 4). Older adults, however, only had slight theta-band motor cortex synchronization toward the end of the 400 ms window (Figures 3, 4). Older adults also showed a slight beta-band (13-35) synchronization in the left posterior parietal cortex before the right mid-extension perturbation (Figure 4). The left posterior parietal cluster had a beta-band synchronization before the left mid-extension perturbations which was followed by a beta desynchronization after the perturbations (Figure 3). Older adults did not show significant posterior parietal spectral fluctuations before or after the perturbation event during left mid-extension (Figure 3).

### 3.2 Baseline Connectivity Analyses

The baseline connectivity analysis revealed significant effects of direction and age group on causal relations between brain and muscle regions (Figure 5). We observed significantly stronger connectivity in the brain-to-muscle direction compared to muscle-to-brain direction (repeated measures ANOVA, F(1,23)=14.9, p*<*0.001). Older adults demonstrated significantly higher connectivity values than younger adults across all conditions (repeated measures ANOVA, F(1,23)=7.5, p=0.012). Task timing showed a marginal main effect on connectivity (repeated measures ANOVA, F(1,23)=3.4, p=0.077).

**Figure 5.**
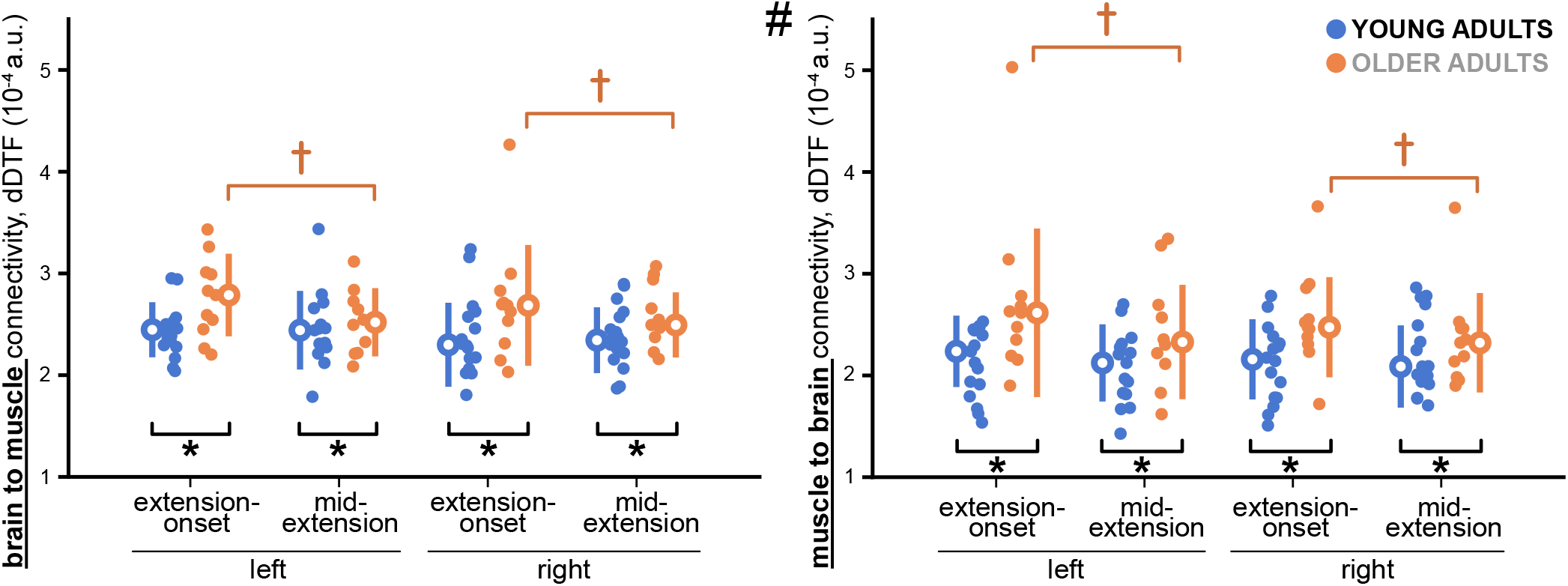
Baseline connectivity prior to the perturbation onset for both perturbation timings, extension- onset and mid-extension. Older adults (orange) had significantly greater baseline connectivity than young adults (blue), as denoted by the *. Older adults also showed significantly greater baseline connectivity at extension-onset compared to mid-extension, as denoted by the †. Brain to muscle connectivity was significantly greater than the muscle to brain connectivity, as denoted by the #.

We also observed a significant task × group interaction (repeated measures ANOVA, F(1,23)=7.5, p=0.012). Post-hoc analysis revealed that older adults showed significantly higher connectivity at extension-onset compared to mid-extension (Bonferroni-corrected, p=0.007), while younger adults showed no significant difference between these time points (p=0.492). Additionally, older adults demonstrated significantly greater connectivity than younger adults specifically at extension-onset (p=0.003), with no group difference at mid-extension (p=0.136). Higher-order interactions (3-way and 4-way) did not reach statistical significance (all p*>*0.05).

### 3.3 Age-related Differences

Event-related corticomuscular connectivity patterns revealed significant age-related differences in neural control strategies during perturbations. To examine these patterns, we analyzed connectivity changes following both left and right perturbations, focusing on left-side muscles to capture non-dominant limb control. Due to the contralateral coupling of the stepper (left leg extends with right arm), the biomechanical roles of muscles switch between perturbation sides. For left perturbations, the driving muscles (shown in green) were posterior deltoid (L.PD), rectus femoris (L.RF), and soleus (L.SO), while the resisting muscles (shown in beige) were anterior deltoid (L.AD), tibialis anterior (L.TA), and semitendinosus (L.ST). For right perturbations, these roles reversed: L.AD, L.TA, and L.ST became the driving muscles, while L.PD, L.RF, and L.SO served as resisting muscles. During left perturbations (Figure 6), older adults demonstrated dense and bidirectional connectivity patterns across all frequency bands, particularly in theta and alpha bands. In theta-band connectivity (Figure 6b), older adults showed strong bidirectional connections from the motor cortex and left posterior parietal cortex (PPC) to nearly all muscles, creating a “broadcast” pattern where cortical regions simultaneously engaged multiple muscles regardless of their biomechanical role. This contrasts with young adults’ sparse, selective theta connectivity, where the anterior cingulate cortex (ACC) showed decreased connectivity to specific muscles (L.PD, a driving muscle, during extension-onset; L.TA, a resisting muscle, during both timings), while supplementary motor areas (SMAs) maintained targeted bidirectional connections primarily with L.AD (a resisting muscle). The distinction between driving and resisting muscles becomes evident in older adults’ connectivity patterns: the motor cortex connected strongly to both driving muscles (L.PD, L.RF, L.SO) and resisting muscles (L.AD, L.TA, L.ST), suggesting a loss of selective muscle control in favor of co-contraction strategies.

**Figure 6.**
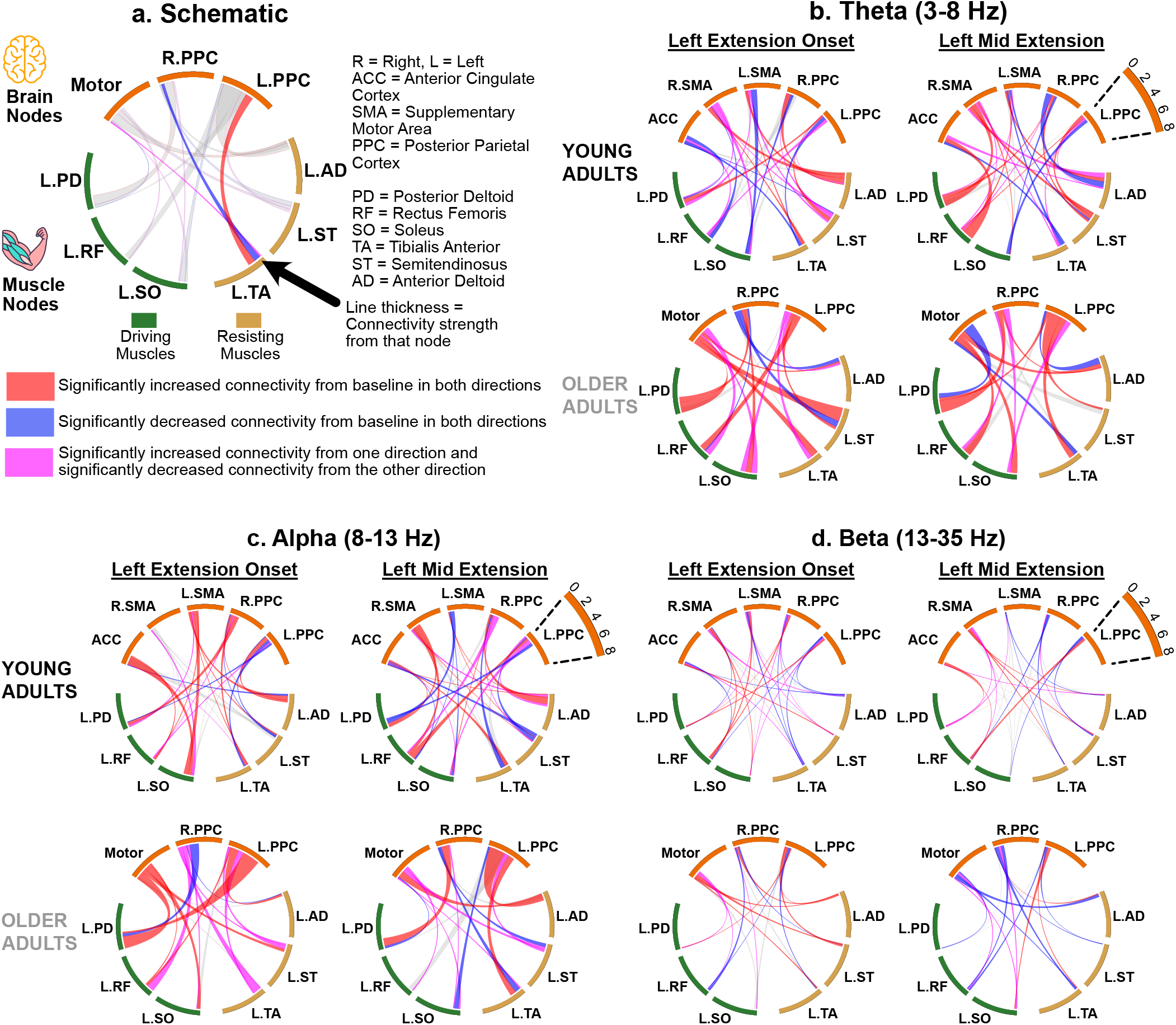
Event related corticomuscular connectivity for the Left Perturbations. (a) Schematic showing an example chord diagram with brain sources (orange) and muscle (driving: green, resisting: beige) labels. All chords are scaled from 0-8e-4. Chord diagrams show average corticomuscular connection strengths during the 400 ms following perturbation onset after baseline subtraction, in (b) theta band, (c) alpha band, and (d) beta band for the left-side perturbations. Only connections that were significantly above or below 0 are shown.

Alpha-band connectivity (Figure 6c) revealed pronounced age-related differences. Older adults exhibited significantly increased bidirectional alpha connections from the motor cortex to both driving muscles (L.PD, L.RF, L.SO) and resisting muscles (L.AD, L.TA, L.ST), with the left PPC connecting to virtually all muscles (*>*4 muscles) during both perturbation timings. This dense alpha connectivity in older adults represents a fundamental reorganization toward constitutive hyperconnectivity, maintaining elevated information flow regardless of specific task demands. Young adults, conversely, showed selective alpha modulation: the ACC decreased connectivity to specific muscles during extension-onset while increasing connectivity to L.RF during mid-extension, demonstrating flexible, context-dependent control. The beta-band connectivity (Figure 6d) further illustrated young adults’ sophisticated control strategies through task-specific SMA lateralization. The left SMA dominated during extension-onset with connections to all muscles except L.SO, then shifted to primarily upper-limb connections during mid-extension, while the right SMA showed the opposite pattern, indicating dynamic hemispheric specialization absent in older adults.

Right perturbations (Figure 7) revealed similar age-related reorganization patterns with notable hemispheric differences. During right perturbations, the muscle roles reversed from left perturbations: L.AD, L.TA, and L.ST served as driving muscles while L.PD, L.RF, and L.SO acted as resisting muscles. In theta-band connectivity (Figure 7b), older adults demonstrated dense, non-selective connectivity from motor cortex and bilateral PPCs to muscles regardless of their biomechanical role. The motor cortex showed strong bidirectional theta connectivity to all muscles during both perturbation timings, contrasting with young adults’ selective ACC and SMA connectivity patterns. The alpha-band responses (Figure 7c) during right perturbations showed particularly strong age-related differences. Older adults exhibited extensive bidirectional alpha connectivity from the right PPC to virtually all muscles during mid-extension, creating an almost complete connectivity network. This hyperconnectivity extended equally to driving muscles (L.AD, L.TA, L.ST) and resisting muscles (L.PD, L.RF, L.SO), suggesting a compensatory strategy that sacrifices selectivity for stability. Young adults maintained their characteristic selective connectivity, with ACC showing targeted modulation and SMAs demonstrating continued lateralization patterns. The beta-band connectivity (Figure 7d) during right perturbations revealed particularly interesting patterns in older adults, with the right PPC showing strong connections primarily to resisting muscles during extension-onset, then shifting to more balanced connectivity during mid-extension, possibly reflecting attempts at impedance control through co-contraction.

**Figure 7.**
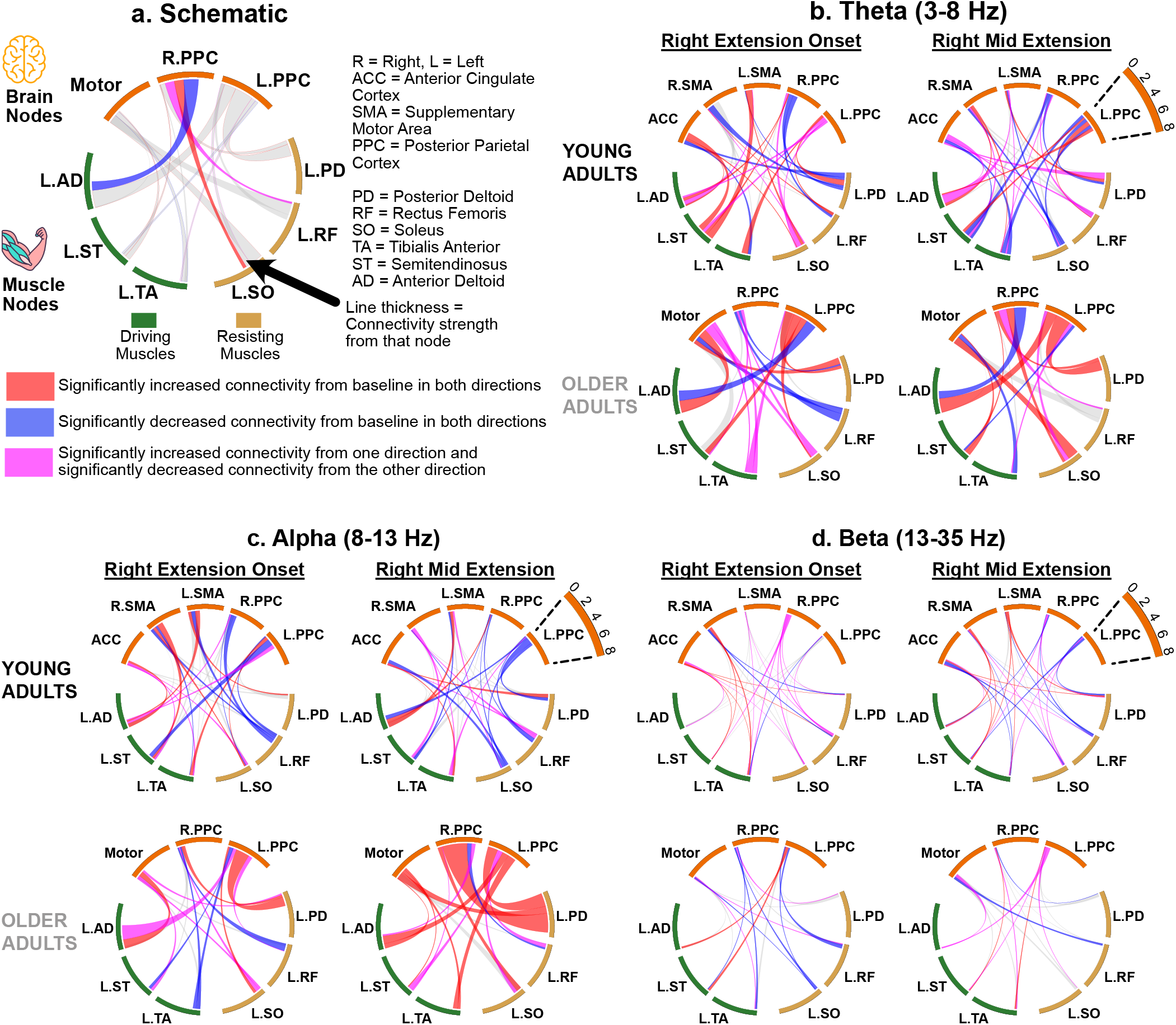
Event related corticomuscular connectivity for the Right Perturbations. (a) Schematic showing an example chord diagram with brain sources (orange) and muscle (driving: green, resisting: beige) labels. All chords are scaled from 0-8e-4. Chord diagrams show average corticomuscular connection strengths during the 400 ms following perturbation onset after baseline subtraction, in (b) theta band, (c) alpha band, and (d) beta band for the right-side perturbations. Only connections that were significantly above or below 0 are shown.

The distinction between driving and resisting muscle connectivity patterns provides crucial insight into age-related motor control changes. Young adults showed task-appropriate connectivity modulation, with selective connections to muscles based on their biomechanical demands. During left perturbations, they modulated connectivity with driving muscles (L.PD, L.RF, L.SO) for force production, while during right perturbations, they shifted to the appropriate driving muscles (L.AD, L.TA, L.ST). Older adults, however, showed equally strong connectivity to both driving and resisting muscles regardless of perturbation side, particularly evident in alpha and theta bands. This non-selective connectivity suggests they employ a co-contraction strategy that trades metabolic efficiency for mechanical stability. The consistency of this pattern across both perturbation types, despite the reversal of muscle roles, indicates a fundamental reorganization of motor control strategy rather than task-specific adaptations.

## 4 Discussion

This study reveals a fundamental architectural reorganization of sensorimotor control with aging, characterized by the loss of error-specific cortical processing and a compensatory shift toward elevated but inflexible corticomuscular connectivity. Young adults had concentrated electrocortical sources in the anterior cingulate, left and right supplementary motor area, and left and right posterior parietal cortex (Figure 2). The anterior cingulate and supplementary motor area clusters consistently showed strong theta-band synchronization after the perturbation event (Figures 3, 4). Older adults demonstrated a critical loss of cortical specialization, with complete absence of anterior cingulate cortex activity and undifferentiated motor regions (Figure 2), suggesting a fundamental degradation of the error-monitoring architecture. The minimal perturbation-locked synchronization in older adults (Figures 3, 4) suggests their cortical networks operate near ceiling, with elevated baseline activity leaving little dynamic range for task-specific modulation.

Directional causal connectivity results reveal young and older adults employing fundamentally divergent control policies. The connectivity analysis shows older adults display significantly stronger baseline corticomuscular connectivity (Figure 5), while the chord diagrams for both left and right perturbations (Figures 6, 7) expose a striking architectural difference: young adults employ sparse, selective connectivity with bidirectional modulation primarily between error-processing regions (ACC, SMAs) and specific muscles, while older adults demonstrate dense, non-selective patterns – a ‘broadcast’ strategy connecting motor and parietal cortices to multiple muscles simultaneously. Critically, the analysis of muscle-specific connectivity reveals that older adults show equally strong connectivity to both driving and resisting muscles, regardless of their biomechanical role in each perturbation. This non-selective connectivity, particularly prominent in theta and alpha bands, suggests a co-contraction strategy that prioritizes mechanical stability. This pattern persists across both left and right perturbations, suggesting not task-specific compensation but rather a fundamental reorganization of motor control architecture. Young adults, conversely, maintain preferential connectivity with driving muscles and show flexible, context-dependent modulation that enables efficient force production while minimizing unnecessary co-contraction. This architectural reorganization represents a trade-off: older adults appear to exchange error-specific processing and dynamic adaptability for constitutive stability, maintaining elevated baseline connectivity as a compensatory strategy while reducing flexible, selective muscle control.

Higher baseline connectivity in older adults, observed without balance demands (Figure 5), aligns with compensatory “overactivation” theories such as the CRUNCH model, where older adults recruit additional neural resources to maintain performance comparable to young adults (Reuter-Lorenz & Cappell, 2008). While previous work demonstrated increased delta and theta CMC during challenging upright stance (Ozdemir et al., 2018), our findings of elevated baseline connectivity during seated stepping suggest that increased cortical engagement in older adults extends beyond reactive postural control, potentially reflecting a fundamental shift in the motor system’s default state.

Our results from anterior cingulate and motor region connectivity support the notion of ACC error monitoring and task-specific lateralization in young adults. Our previous study on recumbent stepping with the same experimental paradigm on young adults showed perturbation-elicited anterior cingulate synchronization in theta band, and task-specific lateralization (Shirazi & Huang, 2021). The anterior cingulate cortex compares the expected motor goals with the sensory inputs (Gehring et al., 2012; Holroyd & Coles, 2002), so we hypothesized that part of the sensory input would come from a subset of muscles, modulating direct causal connectivity to ACC. Analysis of both left and right perturbations revealed complex modulation patterns: we observed bidirectional theta connectivity decreases between ACC and specific muscles (e.g., L.PD during left extension-onset, L.TA during both timings), while other connections showed selective modulation based on task timing (Figures 6b, 7b). This aligns with a previous study showing decreased bidirectional theta connectivity between ACC and right peroneus longus in a “pull” perturbation task during walking (Peterson & Ferris, 2019). The consistency of ACC connectivity patterns across both left and right perturbations suggests that error monitoring operates as a central control mechanism, maintaining selective connections based on functional demands rather than strict anatomical laterality. In addition to bidirectional theta connectivity, bidirectional alpha connectivity patterns (Figures 6c, 7c) revealed task-specific modulation between ACC and both driving and resisting muscles, supporting our hypothesis of cortical control during perturbed locomotion while indicating sophisticated task-specific strategies. A study involving treadmill walking showed significant corticomuscular alpha and beta connectivity from ipsilateral and contralateral motor cortex to lower limb muscles, indicating corticomuscular lateralization (Gwin et al., 2011). Another study found gait-event specific lateralization during overground walking (Artoni et al., 2017a). Our previous study on recumbent stepping (Shirazi & Huang, 2021) showed task-specific lateralization in the SMAs. The connectivity results demonstrate dynamic SMA lateralization across both left and right perturbations: left SMA shows dominant beta connectivity during extension onset which shifts to primarily upper-limb connections during mid-extension, while right SMA exhibits the complementary pattern (Figures 6d, 7d). This bilateral analysis indicates that task-specific CMC lateralization represents an organizational principle in young adults’ motor control that persists across different perturbation conditions.

Older adults showed remarkably strong connectivity patterns in the motor cortex and posterior parietal cortices across both left and right perturbations, revealing profound age-related reorganization of motor control. A study with older adults performing a visually guided walking task found increased theta coherence between motor cortex and tibialis anterior (Malcolm et al., 2021). Other studies have found increased alpha and beta coherence in motor cortex among older adults during challenging motor tasks as well (Johnson & Shinohara, 2012; Kamp et al., 2013; Maidan et al., 2019). We hypothesized that corticomuscular connectivity would increase between motor cortex and the driving muscles (rectus femoris, posterior deltoid). Our bilateral perturbation analysis revealed that older adults showed increased bidirectional connectivity to muscles regardless of their biomechanical role. During left perturbations, they connected equally to driving muscles (L.PD, L.RF, L.SO) and resisting muscles (L.AD, L.TA, L.ST), maintaining this non-selective pattern during right perturbations despite reversed muscle roles. This non-selective connectivity across theta and alpha bands (Figures 6b-c, 7b-c) represents a shift from selective control to a broadcast strategy, engaging multiple muscles simultaneously regardless of biomechanical demands. The pattern was particularly pronounced during right perturbations, where the right PPC showed extensive alpha connectivity to virtually all muscles (Figure 7c), creating a near-complete connectivity network that appears to prioritize stability over selectivity.

The bilateral analysis reveals that older adults’ PPC hyperconnectivity serves a compensatory role in impedance control. Studies with stroke patients had previously indicated the role that the PPC has in joint impedance control during perturbed upper limb tasks (Mutha et al., 2012, 2014). In our study, both left and right PPCs showed strong theta and alpha connectivity to multiple muscles simultaneously (Figures 6b-c, 7b-c), with the connectivity strength often exceeding that of the motor cortex. This bilateral PPC dominance, coupled with equal connectivity to both driving and resisting muscles regardless of perturbation direction, may explain why older adults used fewer muscle pairs with greater co-contraction to drive the stepper (Shirazi & Huang, 2024) – achieving stability through simultaneous muscle activation rather than selective recruitment based on biomechanical demands. This strategy suggests a reorganization from the flexible, error-driven control seen in young adults toward a more constrained, stability-focused approach that maintains behavioral performance despite potentially reduced metabolic efficiency.

While we did not find task-specific lateralization in older adults, they showed specific task-related connectivity patterns and indications of visuomotor processing. Left PPC activity is linked to visual processing during motor tasks, which would suggest an increase in PPC activity in the presence of visual feedback (Mutha et al., 2011, 2014). However, while participants were instructed to follow a visual pacing cue, they did not receive any visual feedback. The presence of strong theta and alpha connectivity at left PPC in older adults may indicate increased reliance on visual processing strategies, even in the absence of explicit visual feedback beyond the pacing cue. This pattern is consistent with less automated and more cognitively demanding neuromuscular control during motor tasks among older adults, as demonstrated in previous studies of cyclic coordination tasks (Heuninckx et al., 2005, 2008). Unlike young adults, we did not find a task-specific lateralization in older adults due to a lack of multiple motor clusters, but older adults showed other task-specific connectivity patterns. For example, motor cortex theta connections and left PPC beta connections were more prominent during mid-extension compared to extension-onset, suggesting task-dependent connectivity modulation in older adults despite the absence of lateralization. Older adults also showed higher baseline connectivity during extension-onset tasks (Figure 5), suggesting a task-dependent change in baseline connectivity. However, recent work found no task-related effects on corticomuscular coherence during visually guided versus unguided stepping (Spedden et al., 2022), suggesting that perturbations may be necessary to elicit task-dependent connectivity modulation.

Limitations of this study include insufficient unperturbed strides for comparative analysis and absence of cross-frequency coherence measures. While dDTF requires at least 100 trials for robust results (Artoni et al., 2017b; Peterson & Ferris, 2019), our study included only *∼*30 catch strides and *∼*50-60 strides in pre/post blocks, preventing reliable unperturbed connectivity analysis. Alternative connectivity methods such as wavelet analyses, or symbolic transfer entropy may be better suited to analyze lower numbers of trials (Arunganesh et al., 2022). The majority of our young adults’ motor cortex responses were at the theta-band, while the muscular activity had high beta and gamma (20+ Hz) frequency activities. This disparity in frequency response might skew dDTF and similar connectivity approaches, which by design look for a causal relation between signals in a similar frequency range (Korzeniewska et al., 2003). Also, the 4 Hz-frequency resolution determined by the dDTF results in just two data points for the theta-band (3-8 Hz) connectivity. Treating different frequency bands as separate signals and fitting multivariate autoregressive models with different orders might result in a stronger representation of theta-band responses in the connectivity analysis. While other studies showed modulation of CMC based on walking speed (Chen et al., 2021; Wei et al., 2021), we did not account for any potential speed-related effects on connectivity since our pacing cue guided the participants to perform the stepping tasks at 60 steps per minute. Having a fixed stepping speed allowed us to observe age-related effects of perturbed recumbent stepping on corticomuscular connectivity in a more controlled setting. Our experimental protocol did not include any self-guided tasks, which limited us to only observing CMC in the presence of visual cue. While self-guided stepping tasks could reveal a more accurate observation on visuomotor processing, absence of this visual cue would likely produce similar connectivity patterns, as pointed out in a study with self-guided and visually guided stepping task (Spedden et al., 2022).

In conclusion, our findings demonstrate fundamental differences in how young and older adults organize corticomuscular connectivity during perturbed recumbent stepping. Young adults employed flexible, error-driven modulation between the ACC, SMAs, and specific muscle groups, enabling selective responses to perturbations. Older adults, lacking distinct ACC activity, relied on diffuse but stable connectivity patterns dominated by motor and posterior parietal cortices, suggesting a strategic shift from reactive error correction to proactive stability maintenance. This reorganization appears adaptive rather than pathological, as older adults maintained comparable behavioral performance through alternative neural strategies. Future investigations should explore the modifiability of this reorganization through targeted interventions, including self-guided stepping, dual-task paradigms, and varied visual feedback conditions (Jensen et al., 2018; Maidan et al., 2019; Malcolm et al., 2021; Seynaeve et al., 2024; Spedden et al., 2022). The distinct connectivity signatures establish perturbed recumbent stepping as a sensitive tool for assessing age-related sensorimotor changes. These normative patterns provide quantitative biomarkers for rehabilitation monitoring and enable personalized interventions targeting specific connectivity deficits – whether restoring youth-like flexibility in early aging or optimizing compensation strategies in advanced aging (Reinkensmeyer et al., 2016).

## Code Availability

The data and code will be available upon request.

## Author contribution

HJH conceived the research and provided funding. SYS and HJH designed the experiment, collected data, performed pre- and post-processing. SMT reanalyzed the connectivity results, regenerated the figures and performed statistical analysis. SYS, HJH, and SMT made the figures and wrote the manuscript.

## Acknowledgments

This work was partially supported by the National Institute on Aging of the National Institutes of Health under R01AG054621.

